# Genome compartmentalization predates species divergence in the plant pathogen genus *Zymoseptoria*

**DOI:** 10.1101/864561

**Authors:** Alice Feurtey, Cécile Lorrain, Daniel Croll, Christoph Eschenbrenner, Michael Freitag, Michael Habig, Janine Haueisen, Mareike Möller, Klaas Schotanus, Eva H. Stukenbrock

## Abstract

**Background:** Antagonistic co-evolution can drive rapid adaptation in pathogens and shape genome architecture. Comparative genome analyses of several fungal pathogens revealed highly variable genomes, for many species characterized by specific repeat-rich genome compartments with exceptionally high sequence variability. Dynamic genome architecture may enable fast adaptation to host genetics. The wheat pathogen *Zymoseptoria tritici* with its highly variable genome and has emerged as a model organism to study the genomic evolution of plant pathogens. Here, we compared genomes of *Z. tritici* isolates and genomes of sister species infecting wild grasses to address the evolution of genome composition and structure.

**Results:** Using long-read technology, we sequenced and assembled genomes of *Z. ardabiliae*, *Z. brevis*, *Z. pseudotritici* and *Z. passerinii*, together with two isolates of *Z. tritici*. We report a high extent of genome collinearity among *Zymoseptoria* species and high conservation of genomic, transcriptomic and epigenomic signatures of compartmentalization. We identify high gene content variability both within and between species. In addition, such variability is mainly limited to the accessory chromosomes and accessory compartments. Despite strong host specificity and non-overlapping host-range between species, effectors are mainly shared among *Zymoseptoria* species, yet exhibiting a high level of presence-absence polymorphism within *Z. tritici*. Using *in planta* transcriptomic data from *Z. tritici*, we suggest different roles for the shared orthologs and for the accessory genes during infection of their hosts.

**Conclusion:** Despite previous reports of high genomic plasticity in *Z. tritici*, we describe here a high level of conservation in genomic, epigenomic and transcriptomic signatures of genome architecture and compartmentalization across the genus *Zymoseptoria*. The compartmentalized genome may reflect purifying selection to retain a functional core genome and relaxed selection on the accessory genome allowing a higher extent of polymorphism.

## Introduction

Co-evolution between plants and pathogens can drive rapid evolution of genes involved in antagonistic interaction [1]. In filamentous plant pathogens, rapid evolution may be fueled by highly dynamic genome architecture involving repeat-rich compartments such as gene-sparse islands of repetitive DNA and accessory chromosomes [2, 3]. These compartments can show a high plasticity revealed by a high extent of gene and/or chromosome presence-absence variation and structural variants, such as inversions, insertions and deletions [4, 5]. Several plant pathogenic fungi have isolate specific chromosomes, so-called accessory chromosomes.

Accessory chromosomes are characterized by intra-species presence-absence polymorphism, low gene density, an enrichment of repetitive sequences and, in some species, a different histone methylation pattern [6, 7]. It has been shown that accessory chromosomes encode genes involved in virulence such as in the species *Fusarium solani*, *Fusarium oxysporum* and *Leptosphaeria maculans* [8–11]. Little is known about the evolutionary origin of accessory chromosomes although experimental evidence from the asexual species *F. oxysporum* shows that accessory chromosomes may be acquired horizontally as chromosomes can be transferred between distinct isolates by hyphal fusion [10]. Through such transfers, virulence determinants may be exchanged between clonal lineages as accessory chromosomes in this species were shown to encode host specific virulence determinants and transcription factors regulating their expression [12].

Genes involved in plant-pathogen interactions may diversify at a higher rate in repeat-rich genome compartments and thereby evolve new virulence specificity faster [3]. These genes encode secreted proteins, so-called effectors [1]. Most known effectors target diverse cellular compartments and molecular pathways, including immune response-related pathways [13, 14]. Genes encoding Carbohydrate-active enzymes (CAZymes) have also been associated to the pathogenic lifestyle of fungal plant pathogens, particularly through their role in plant-cell wall degradation [15]. Thus, some secreted CAZymes may be essential from the early infection stage, like penetration of plant tissue, to later stages such as the necrotrophic phase where the pathogen feed from dead plant tissue [16]. Likewise, secondary metabolites are known to be involved in plant infection and contribute to virulence and the interaction with other plant-associated microorganisms [17, 18]. Many of these genes can be predicted either according to their composition and known protein domains or through machine learning methods [19]. Thereby, in-depth genome annotations have proven important to predict and compare the content of pathogenicity-related genes in plant pathogens, as well as their genomic localization for example in rapidly evolving genome compartments.

The ascomycete pathogen *Zymoseptoria tritici* has emerged as a model organism in evolutionary genomics of pathogens. This species originated in the Fertile Crescent during the domestication of its host, wheat [20]. Closely related species of *Z. tritici* have been collected from wild grasses in the Middle East providing an excellent resource for comparative genome analyses of closely related and recently diverged pathogen species. Comparative analyses of genome organization and gene content within and among *Zymoseptoria* species have previously revealed a wide distribution of accessory chromosomes and dynamic gene content [21, 22]. The haploid genome of the reference isolate IPO323 comprises thirteen core and eight accessory chromosomes [23]. Some of these accessory chromosomes encode traits that impact virulence of the fungus, however no gene encoded on an accessory chromosome has so far been described as a virulence or avirulence determinant [24–29]. Interestingly, the accessory chromosomes in *Z. tritici* show a low transcriptional activity *in vitro* as well as *in planta* [30, 31]. This suppression of gene expression correlates with an enrichment of heterochromatin associated with the histone modification H3K27me3 on the accessory chromosomes [6, 32].

In the reference isolate IPO323, the accessory chromosomes comprise more than 12% of the entire genome assembly. To which extent such a high amount of accessory DNA is also found in genomes of other members of the *Zymoseptoria* genus has so far been unknown due to the lack of high-quality genome assemblies and large scale population sequencing. Assemblies based on short-read data failed to recover complete sequence of accessory chromosomes and “orphan regions” due to their high repeat content [22]. The asset of genome assemblies based on long-read sequencing was demonstrated in a detailed genome comparison of four *Z. tritici* isolates sequenced with PacBio long-read sequencing [28]. Comparison of the high-quality chromosome assemblies revealed the occurrence of “orphan regions” enriched with transposable elements and encoding putative virulence-related genes [33].

In this study, we investigate the genomic architecture and variability among five *Zymoseptoria* genomes. Beside presenting a new and significantly improved resource for future genomic studies of these fungal pathogens, we specifically ask: 1) how conserved is the genome architecture among *Zymoseptoria* species, 2) can we identify accessory compartments in other *Zymoseptoria* isolates and 3) to which extent does variation in genome architecture reflect variation in gene content.

To answer these questions, we used high-quality assemblies based on long-read sequence data and new gene predictions in two isolates of *Z. tritici* (Zt05 and Zt10) and one isolate of each of the sister species, *Z. ardabiliae, Z. brevis, Z. passerinii,* and *Z. pseudotritici.* We explore the core and non-core genome architecture of *Zymoseptoria* spp. combining genomic data with transcriptome and histone methylation data and relate this to core and accessory genome compartments. Furthermore, we compare the distribution of orthologous and non-orthologous genes in the *Zymoseptoria* genomes and one additional Dothideomycete species. Our analyses reveal an overall conserved genome architecture characterized by gene-rich core compartments and accessory compartments enriched in species-specific genes. Finally, we report an exceptionally high extent of variation in presence-absence of protein coding genes in a eukaryote genome.

## Results

### New de novo assemblies using long-read sequencing for six *Zymoseptoria* spp

We sequenced and assembled the genome of the reference isolates of *Z. ardabiliae*, *Z. brevis*, *Z. pseudotritici* and *Z. passerinii* and the genomes of two *Z. tritici* isolates sampled in Denmark and Iran. The obtained contigs were filtered based on base-quality confidence and read depth to ensure high quality of the final assemblies (see Methods). This filter removed a high number of contigs (between 17% and 58% of the total), but little overall length (between 0.4 and 2.6% of the total assemblies), indicating that most of the excluded contigs were of small size. The best assemblies were of the two *Z. tritici* isolates comprising 19 and 30 contigs and the most fragmented was of *Z. passerinii* comprising 103 contigs (Figure 1). The resulting assembly lengths ranged from 38.1 Mb for *Z. ardabiliae* to 41.6 Mb for *Z. brevis*, which is comparable to the reference assembly length of *Z. tritici* (39.7 Mb) but slightly larger than previous short-read based assemblies (Table 1; previous assemblies ranged from 31.5 *Z. ardabiliae* to 32.7 for *Z. pseudotritici* [22, 23]. The assembly of the Iranian *Z. tritici* isolate Zt10 has telomeric repeats at the end of all contigs, indicating that each chromosome is completely assembled, comprising six accessory and thirteen core chromosomes. The assemblies for the Danish *Z. tritici* isolate (Zt05), *Z. brevis* (Zb87) and *Z. pseudotritici* (Zp13) contained, respectively, twelve, nine and five fully assembled chromosomes including both core and accessory chromosomes (Figure S1). The assemblies of the *Z. ardabiliae* (Za17) and *Z. passerinii* (Zpa63) genomes included no fully assembled chromosomes, but twelve and ten contigs respectively with telomeres at one of the ends (Table 1; Figure S1).

**Figure 1:**
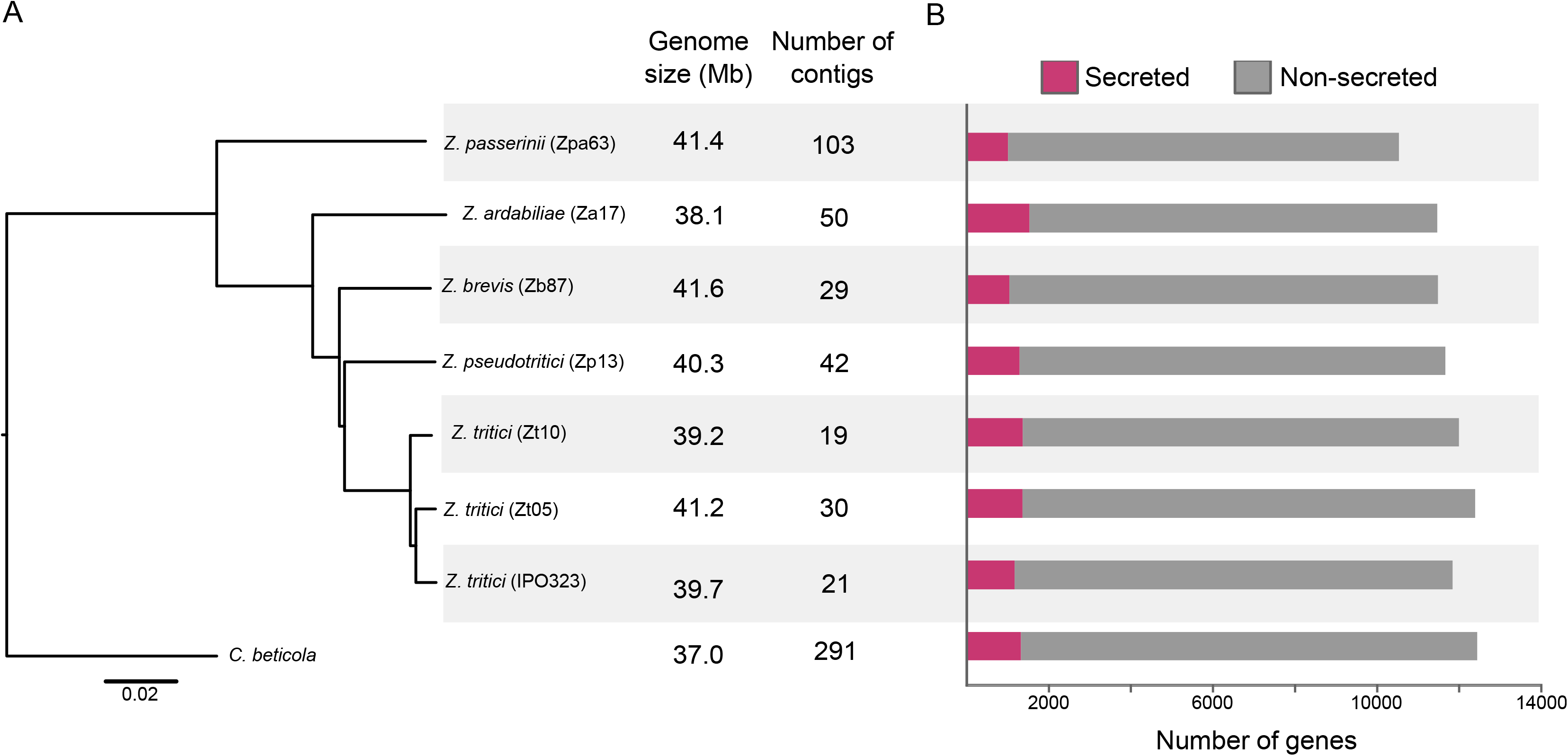
Whole-genome phylogeny of *Zymoseptoria* spp. and basic statistics for the assemblies and gene predictions. Tree based on the distance matrix generated from the whole-genome assemblies using the software andi (Haubold et al. 2015). The bar plots represent the number of genes coding for secreted proteins (pink) and non-secreted proteins (grey) for each genome.

**Table 1:**
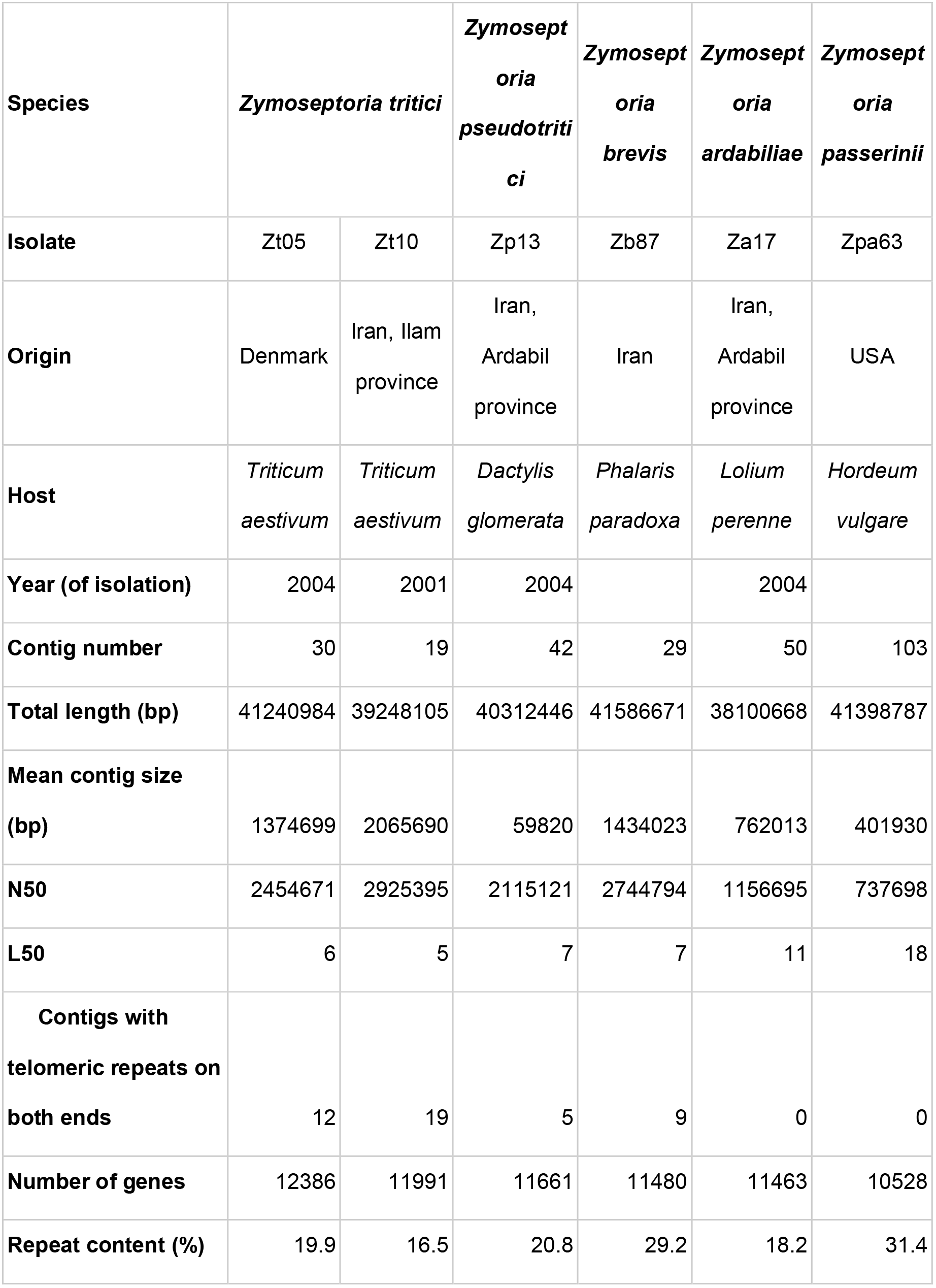
Metrics of genome assemblies and annotation

| **Species** | ***Zymoseptoria tritici*** |  | ***Zymoseptoria pseudotritici*** | ***Zymoseptoria brevis*** | ***Zymoseptoria ardabiliae*** | ***Zymoseptoria passerinii*** |
| --- | --- | --- | --- | --- | --- | --- |
| **Isolate** | Zt05 | Zt10 | Zp13 | Zb87 | Za17 | Zpa63 |
| **Origin** | Denmark | Iran, Ilam province | Iran, Ardabil province | Iran | Iran, Ardabil province | USA |
| **Host** | *Triticum aestivum* | *Triticum aestivum* | *Dactylis glomerata* | *Phalaris paradoxa* | *Lolium perenne* | *Hordeum vulgare* |
| **Year (of isolation)** | 2004 | 2001 | 2004 |  | 2004 |  |
| **Contig number** | 30 | 19 | 42 | 29 | 50 | 103 |
| **Total length (bp)** | 41240984 | 39248105 | 40312446 | 41586671 | 38100668 | 41398787 |
| **Mean contig size (bp)** | 1374699 | 2065690 | 59820 | 1434023 | 762013 | 401930 |
| **N50** | 2454671 | 2925395 | 2115121 | 2744794 | 1156695 | 737698 |
| **L50** | 6 | 5 | 7 | 7 | 11 | 18 |
| **Contigs with telomeric repeats on both ends** | 12 | 19 | 5 | 9 | 0 | 0 |
| **Number of genes** | 12386 | 11991 | 11661 | 11480 | 11463 | 10528 |
| **Repeat content (%)** | 19.9 | 16.5 | 20.8 | 29.2 | 18.2 | 31.4 |

The transcriptome-based gene predictions for these new assemblies include between 10,528 and 12,386 protein-coding genes (Table S1). This range is consistent with the annotation of the reference genome IPO323 reporting 11,839 protein-coding genes [22]. We used Benchmarking Universal Single-Copy Orthologs (BUSCO) from the lineage dataset *Pezizomycotina* to evaluate the completeness of the assemblies and gene predictions (Waterhouse et al. 2018). The proportion of complete BUSCOs were as follow Zt10: 98.7%, Zt05: 98.4%, Zp13: 98.5%, Zb87: 97.0%, Za17: 98.2% and Zpa63: 97.5%. These values are comparable to the one obtained for the reference genome of *Z. tritici* (97.8%). The assessment of gene content completeness (see Methods) indicates that, despite more fragmented assemblies of *Z. ardabiliae* and *Z. passerinii*, the genomes are complete in terms of gene content and that the unassembled fragments are more likely to comprise repeats and not protein coding genes.

Based on the whole-genome sequences and the predicted genes we reconstructed the phylogeny of the *Zymoseptoria* genus using the publicly available genome of *Cercospora beticola* as an outgroup [34, 35]. For both trees, the phylogenetic relationship of the *Zymoseptoria* species is in accordance with previously published phylogeny based on seven loci sequenced in multiples isolates (Figure 1) [21].

### Genomes of *Zymoseptoria* spp. comprise accessory chromosomes and compartments but show overall high synteny

Next we addressed the extent of co-linearity of the *Zymoseptoria* genomes. Using coordinates of orthologous genes, we were able to reveal a high extent of synteny conservation among the five *Zymoseptoria* species and between the three isolates of *Z. tritici*, as depicted in Figures 2 and S2. Based on this high extent of synteny and the prediction of telomeric repeats, we identified the correspondence of chromosomes between the reference genome of *Z. tritici* IPO323 and the other *Zymoseptoria* genomes (Figure 2 and S2). *Z. brevis* and *Z. pseudotritici* share a near perfect synteny in their core chromosomes, however, when compared to *Z. tritici*, *Z. brevis* and *Z. pseudotritici* have two large-scale inversions comprising roughly ~900 kb and ~1.2 Mb of chromosomes 2 and 6, respectively (Figure 2, S2 and S3). Based on the phylogeny in Figure 1, it is likely that these two events occurred after the divergence of *Z. tritici* from *Z. brevis* and *Z. pseudotritici*. Overall, we observe a higher extent of synteny conservation between *Z. brevis* and *Z. pseudotritici* compared to *Z. tritici* IPO323 (Figure 2; S2 and S3).

**Figure 2:**
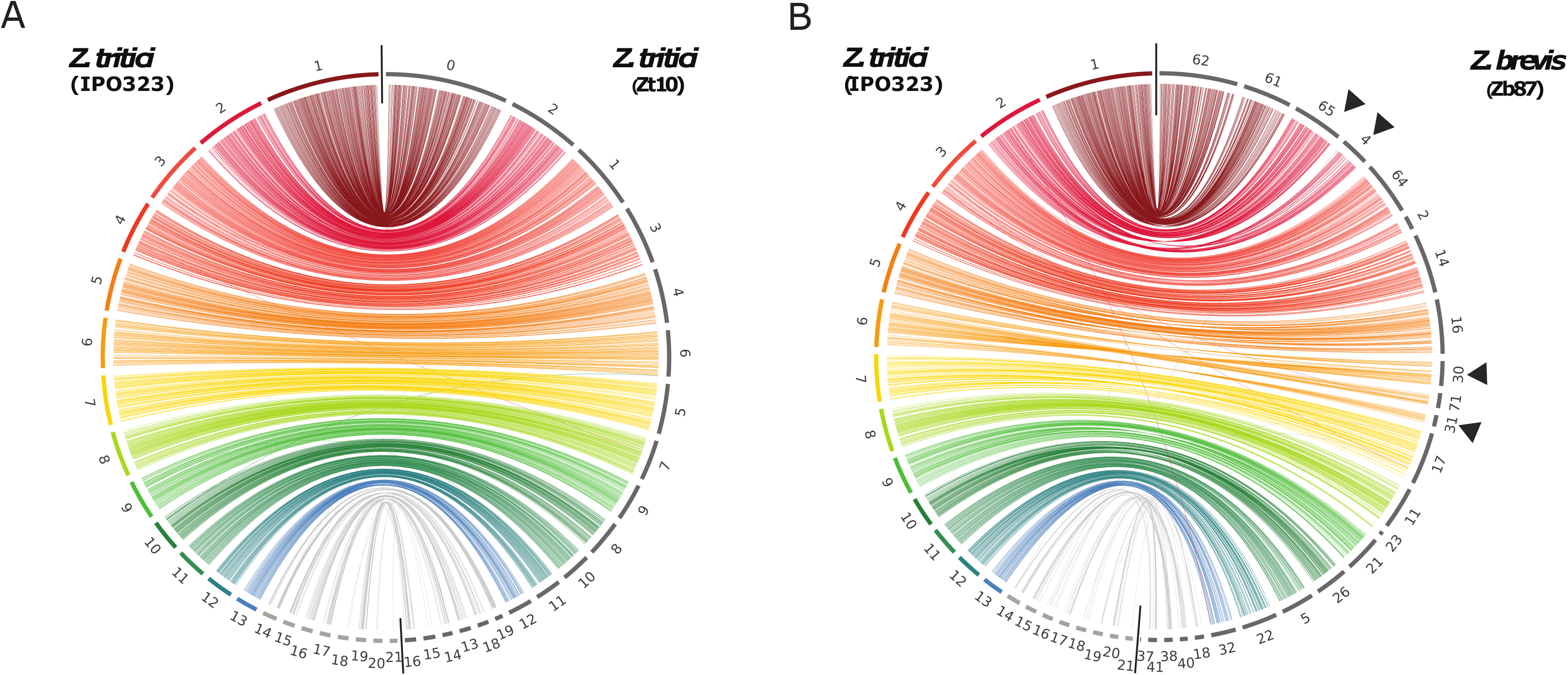
Intra- and inter-species synteny conservation in *Zymoseptoria* genus. A) Intra-species synteny between the reference genome of *Z. tritici* and the genome of the Iranian *Z. tritici* isolate Zt10. Each color represents a different chromosome as defined in the reference *Z. tritici* genome, except for accessory chromosomes, which are in grey. The links represent orthologous genes. B) Inter-species synteny between the reference genome of *Z. tritici* and the genome of *Z. brevis*. The arrows represent the large-scale inversions identified between the genomes of these two species.

In *Z. tritici*, core and accessory compartments have very distinct genomic features. It was previously shown that hallmarks of accessory regions in the reference isolate IPO323 include lower gene density, lower levels of H3K4 methylation levels and lower gene expression [6, 31]. In the reference genome of IPO323, compartments with these genomic and epigenomic hallmarks represent either accessory chromosomes or specific regions of the core chromosomes. Here we find that the specific accessory hallmarks including low gene density, low expression, low H3K4me2 methylation and significant enrichment of species-specific genes (see description below) on the non-core contigs are found in genomic compartments throughout the genus (Table S3, Figures 3 and S4).

**Figure 3:**
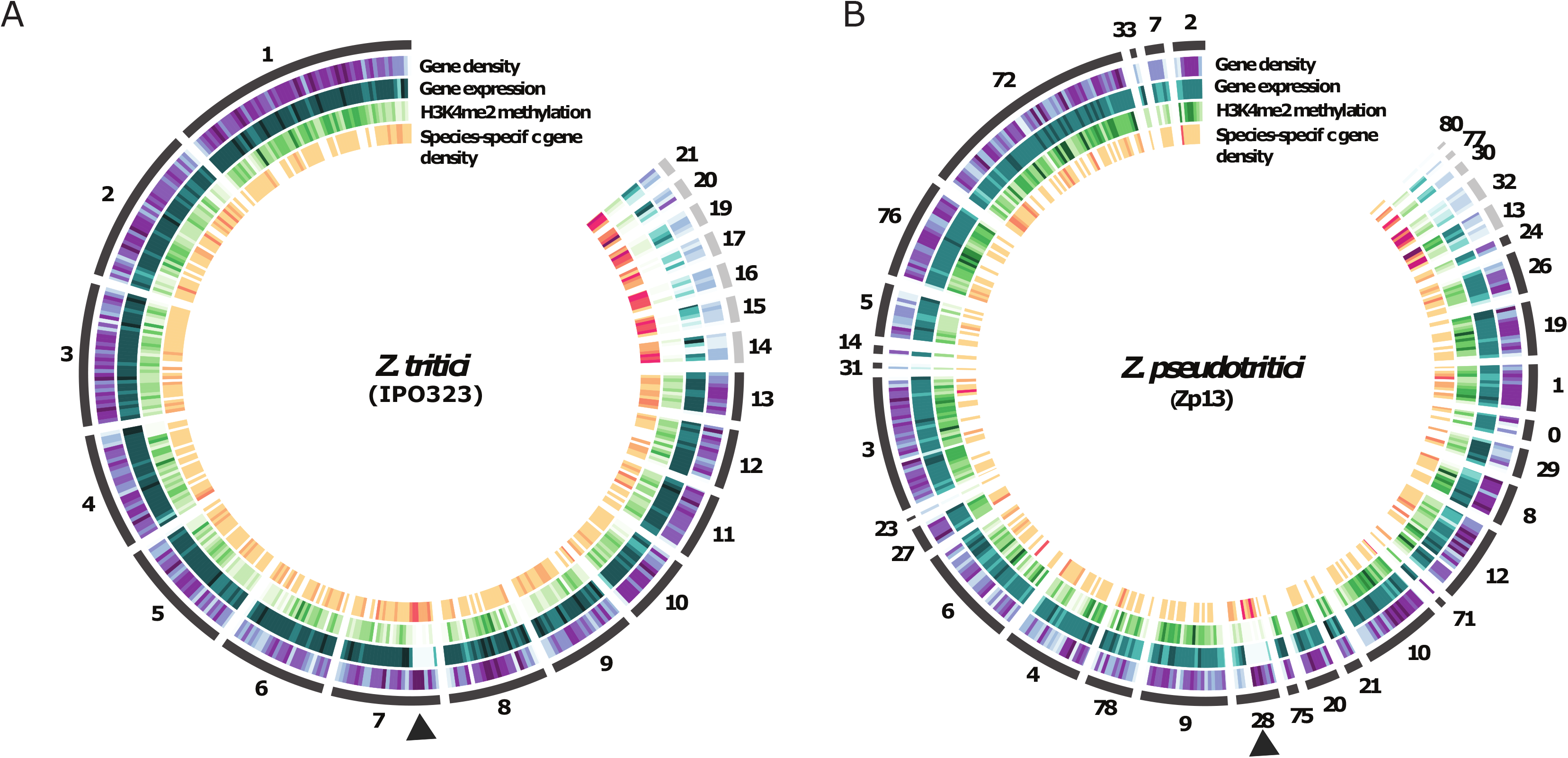
Genome architecture of the reference genome *Z. tritici* IPO323 (A) and *Z. pseudotritici* Zp13 (B). The segments constituting the first circle represents the chromosomes of IPO323 (A) and contigs of Zp13 (B) ordered according to chromosome numbers of the reference genome. Tracks from the outside to the inside are heatmaps representing respectively: gene density along chromosomes/contigs; gene expression in vitro (TPM); H3K4me2 levels in vitro and species-specific gene density. The arrows indicate the location of the region on chromosome 7 (and the corresponding syntenic region in Z.pseudotritici) displaying accessory-like genomic and regulatory hallmarks.

In the reference *Z. tritici* strain, the specific “accessory-like” pattern includes a particular region of the core chromosome 7 of ~0.6 Mb (Figure 3A) [6]. This particularly large accessory marked region is observed in several *Zymoseptoria* spp (Figure 3B; S4). We identify ~0.7 Mb of the contig 28 in *Z. pseudotritici* and ~0.6 Mb of the contig 17 for *Z. brevis*, both corresponding to chromosome 7 of *Z. tritici*, which share the same hallmarks of accessory chromosomes (Figures 3 and S4, Table S3). For the two remaining sister species, the fragmentation of the assembly does not allow the identification of such pattern although there is an indication of a similar effect on gene expression and species-specific gene enrichment on contig 19 of *Z. ardabiliae* which corresponds to a fragment of chromosome 7 in IPO323 (Figure S5).

We also identified other regions enriched in isolate-specific genes, thus defining orphan loci in the core chromosomes of both Zt10 and Zt05 (Figures 3 and S4). We observe a region of ~0.2 Mb of contig 1 in the Iranian isolate Zt10 corresponding to the core chromosome 3 in the IPO323 genome with high content of isolate-specific genes (Figure S4; table S3). We furthermore identified small segments with species-specific genes on the core chromosomes of the wild-grass infecting sister species including a ~0.3 Mb region of the contig 26 in *Z. brevis* and ~0.1 Mb of contig 30. Overall, we show that genome compartmentalization in core and accessory regions is an ancestral and shared trait among the *Zymoseptoria* species. This phenomenon generates highly variable compartments and defined loci that deviate from genome averages in terms of gene content, sequence composition and synteny conservation.

### Variable repertoires of effector candidate genes

To obtain gene annotations for the *Zymoseptoria* genome assemblies, we established a custom pipeline adapted from Lorrain and co-workers (Figure S5) [36]. Briefly, we use the consensus of three methods to predict gene product localization, then extract secreted proteins to further identify predicted effectors. This detailed functional annotation provided a catalog of predicted gene functions and cellular localizations (Figure S6). For each genome, a large proportion of genes could not be assigned to a protein function and lacked protein domains. 49.6% of genes in *Z. tritici* (N = 5953) and up to 71.8% of genes in *Z. pseudotritici* (N = 8373) lack a predicated function (Figure S6A). A relatively consistent number of genes are predicted for each functional category among *Zymoseptoria* spp. (Figure S6A and B). Likewise, the numbers of gene products predicted to belong to the different subcellular localizations are very similar (Figure S6C) across the whole genus, including secreted proteins. The difference between the minimal and maximal gene number fpr the different categories of subcellular localizations do not exceed 1.6X between species (Figure S6C). Overall, secretomes range from 7% of the genes predicted in *Z. passerinii* (N=828) to 11% of genes in *Z. ardabiliae* (N=1328 genes, Figure S6B).

We further investigated the number and distribution of genes predicted to encode proteins with a pathogenicity-related function, such as secondary metabolites, CAZymes and effector candidates (Figures S1 and S6B). Genes involved in the synthesis of secondary metabolites are typically organized in clusters, with genes participating in the same biosynthetic pathway grouping together at a genomic locus (Shi-Kunne et al. 2019). The number of biosynthetic gene clusters (BGC) ranges from 25 in *Z. ardabiliae* and *Z. passerinii* to 33 in the IPO323 reference genome and includes from 305 to 471 predicted genes (Figure S1). The only BGC identified in a non-core contig is a non-ribosomal peptide synthetase BGC found on the contig 38 of *Z. brevis* which has no orthologous cluster detected in any of the other *Zymoseptoria* genomes (Figure S1). We identified between 454 and 515 CAZyme genes in the *Zymoseptoria* species. Both BGCs and CAZymes are almost exclusively found on the core chromosomes (Figure S1). The only exceptions are a CAZyme encoding gene found on chromosome 14 in *Z. tritici* IPO323 and Zt05, and a CAZyme encoding gene on the putative accessory contig 38 of *Z. brevis* (Figure S1). These two genes encode for a beta-glucosidase and a carboxylic-ester hydrolase, respectively.

In contrast to the high conservation of CAZyme and BGC gene content among the *Zymoseptoria* genomes, we find that predicted effector genes exhibit a large variation in gene numbers between genomes (Figure S6B). In fact, the predicted effector gene repertoire of in *Z. ardabiliae* (N=637) is three times as high compared to *Z. brevis* (N=206). Interestingly, the three *Z. tritici* isolates also vary considerably in their predicted effector repertoires. The reference isolate IPO323 has a reduced set of effector genes (N=274) compared to Zt05 and Zt10 that encode approximately 30% more effector genes (N=417 and N=403, respectively, Figure S6B). Despite the high variability, the effector genes are mostly located on core chromosomes and none of the five *Zymoseptoria* species have more than ten effector genes located on accessory chromosomes (Figure S1).

### The accessory genes of *Z. tritici* are shared with the closely related wild-grass infecting species

To further characterize variation in gene content among the five *Zymoseptoria* species, we identified orthologous genes (i.e. orthogroups) from the gene predictions. We categorized 22341 gene orthogroups identified in the seven *Zymoseptoria* genomes and in *C. beticola* according to their distribution among fungal genomes (Figure 4A). The core orthogroups, which are genes present in all eight genomes, represent around 30% of all orthogroups (N = 6698). The *genus-specific* orthogroups, shared between several *Zymoseptoria* spp. but not found in the *C. beticola* genome, represent 45% of the orthogroups (N = 9955; ranging from 2066 to 3212 per species). Among the *genus-specific* orthogroups, 1100 are found in all *Zymoseptoria* genomes (Figure 4A), whereas all others show presence-absence polymorphisms within the genus. A total of 2476 *species-specific* orthogroups (ranging from 552 to 1191 per species) are found only in individual species. Among the *species-specific* genes, 205 orthogroups (N_genes_ = 414 to 562) are found in all three *Z. tritici* genomes while the *isolate-specific* genes in *Z. tritici* represent 391 (Zt10) to 792 (IPO323) genes.

**Figure 4:**
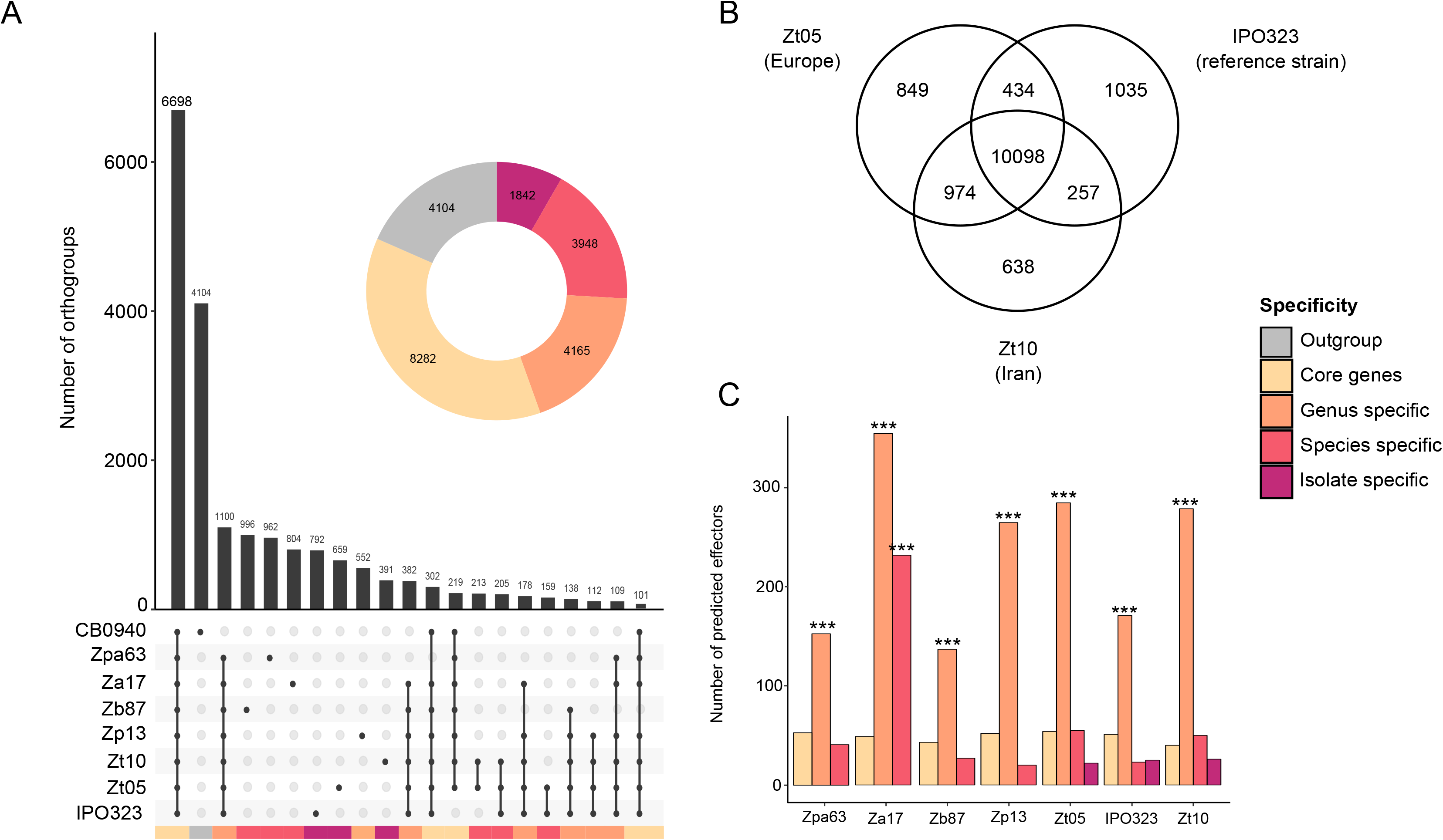
Orthogroups and functional gene categories in *Zymoseptoria* spp. genomes. A) Orthogroups shared by the reference *Z. tritici* genome, our new *Zymoseptoria* assemblies and the outgroup genome of *C. beticola*. Only intersects higher than 100 are displayed on the upset plot. The doughnut plot summarizes the number of orthologs grouped by larger categories: specific to some isolates, to a species or shared by all. The colored bars under the upset plot link each intersect to its corresponding category in the doughnut plot. B) Venn diagram representing the genes shared by the three isolates of *Z. tritici*. C) The only gene category found to be overrepresented in any of the specificity categories - other than unknown function genes - are effectors. Effectors genes are overrepresented in the genus-specific genes and in *Z. ardabiliae* specific genes (*** represent Fisher exact test p-value < 0.05).

Comparing the three *Z. tritici* isolates independently from the other species, we observe extensive gene presence-absence polymorphisms between the three isolates: 1540 orthogroups are identified in only two strains and 2522 are found in only one (Figure 4B). The number of genes showing presence-absence variation is striking compared to the 10098 core genes in *Z. tritici* as these genes comprise almost 30% of all predicted genes. Interestingly, we show that the number of orthogroups detected as *isolate-specific* is much larger when the comparison includes only members of the same species than when the other species are included (1035, 849 and 638 vs 792, 659 and 391 genes for IPO323, Zt05 and Zt10 respectively; Figures 4A and B). This indicates that a large part of the accessory gene content in *Z. tritici* is shared among the sister species, and highlight the importance of including sister species when establishing core and accessory gene content.

Interestingly, we show that effectors are enriched among the *genus-specific* genes but not among the *species-specific* or *isolate-specific* gene categories (with the exception of *Z. ardabiliae*). Fifty-six percent of effectors in *Z. ardabiliae* and up to 78% of effectors in *Z. pseudotritici* are shared with at least one of the other five *Zymoseptoria* species (Figure 4C). Indeed, 427 effector orthogroups are found in at least two genomes. However, only 47 (10% of the total effector orthogroups N= 474) are found in all seven *Zymoseptoria* genomes. Among the effectors shared by *Z. tritici,* and at least one other *Zymoseptoria* species, 32% (N = 112 of 352) are present in all three *Z. tritici* isolates while 68% (N = 240 of 352) show presence-absence polymorphisms in at least one of the three isolates. These results indicate that the majority of these shared effectors is actually accessory (i.e. presence-absence polymorphism) in *Z. tritici*.

### Among *in planta* differentially expressed genes, species-specific are more expressed than core genes

Finally, we addressed the functional relevance of accessory and orphan genes in *Z. tritici* by analyzing gene expression patterns. We used previously published transcriptome data of three *Z. tritici* isolates [30] and focused on gene expression of the above-defined categories (*core genes, genus-specific, species-specific, and isolate-specific*). We sorted *in planta* expression data into two different infection phases: the biotrophic phase and the necrotrophic phase, a separation supported by principal component analysis of normalized DESeq2 counts (Figure S7). We compared expression levels by mapping RNA-seq reads to the genomes of IPO323, Zt05 and Zt10, using normalized read mappings to transcript per million. We tested differences among gene categories using pairwise comparisons with a Kruskal-Wallis test (Figure 5). Overall, we find that gene expression of the *species-specific* and *isolate-specific* genes is significantly lower in IPO323 and Zt10, but not in Zt05 (Kruskal-Wallis p-value < 0.05). *Species-specific* and *isolate-specific* gene median expression ranges from 3.2 to 5.6 TPM in IPO323 and Zt10 while median expression of core genes is 12.1 and 10.9, respectively. The Zt05 expression profile does not follow the same trend: the *core genes* are the lowest expressed gene category (8.9 median TPM), while *genus*-; *species*- and *isolate-specific* genes showed higher transcription levels (12.0; 14.4 and 13.5 median TPM respectively, Kruskal-Wallis p-value < 0.05).

**Figure 5:**
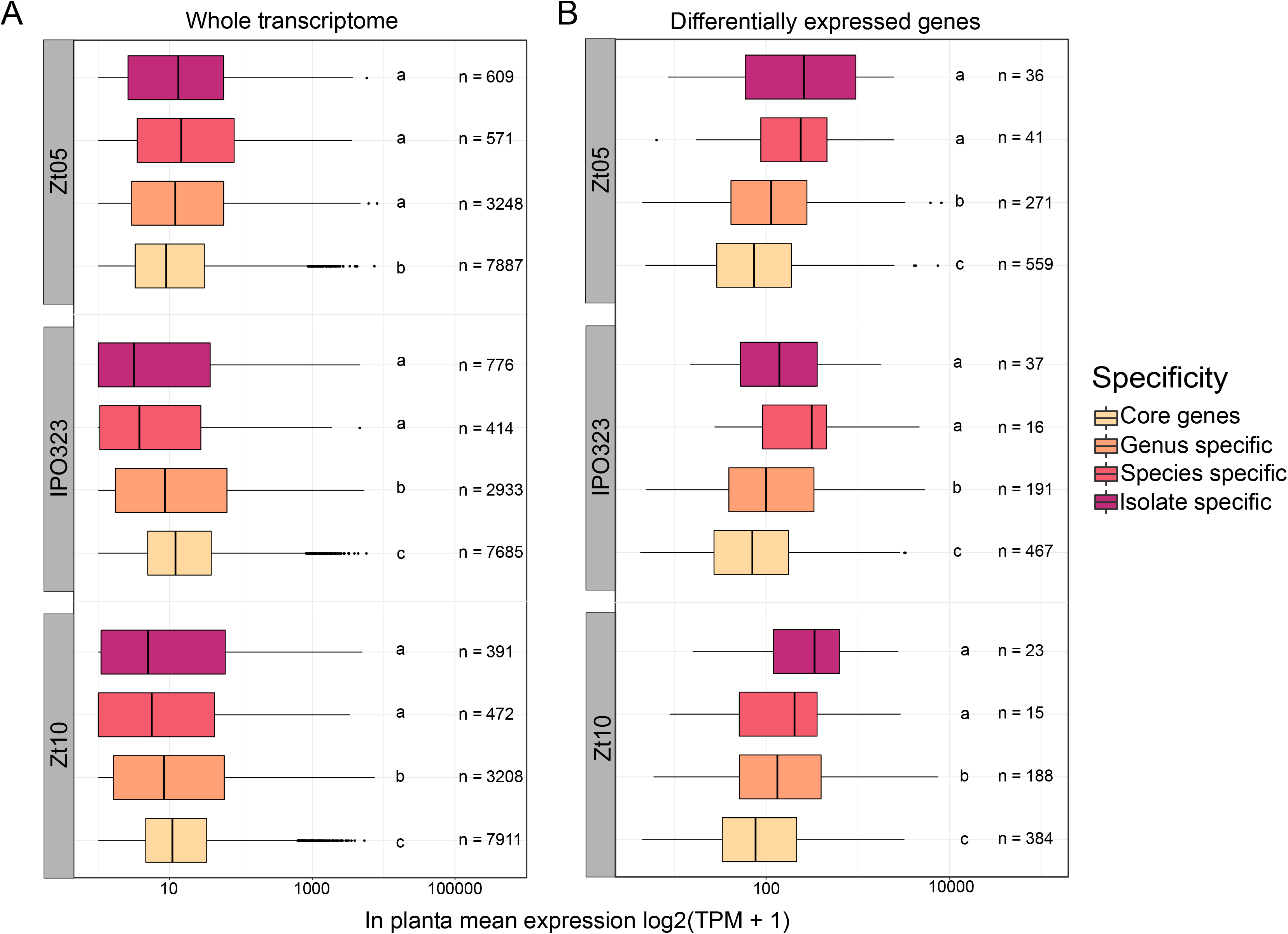
Expression of genes belonging to different specificity levels in the *Zymoseptoria* pangenome. The boxplots represent the expression levels in both biotrophic and necrotrophic phase in transcript per million (TPM) for A) the whole transcriptome of *Z. tritici* isolates and B) in planta differentially expressed genes identified by DESeq2. Comparisons are performed by Krustal-Wallis test, different letters represent p-value < 0.05.

In contrast, we observe a significantly higher expression of the *species-specific* and *isolate-specific* genes for all three isolates (Table S2; Kruskal-Wallis p-value < 0.05) when comparing the expression of genes that are differentially expressed (DEGs; DESeq2 p-adjusted < 0.05) between the biotrophic and necrotrophic phases. *Species-specific* and *isolate-specific* DEGs are higher expressed *in planta* than the *core* and *genus-specific* genes (Figure 5B; Kruskal-Wallis p-value < 0.05). The expression pattern of DEGs with different levels of specificity present a consistent pattern in all three isolates (Table S2). Overall, this comparison reveals a potential functional relevance of accessory genes which are up-regulated during infection of *Z. tritici*.

## Discussion

In this study we present a new resource of high-quality whole genome assemblies and gene annotations for the species *Z. ardabiliae, Z. brevis, Z. passerinii, Z. pseudotritici,* and two *Z. tritici* isolates. This new dataset provides a valuable resource for detailed analyses of genome architecture and evolutionary trajectories in this group of plant pathogens. Here, we conduct some first comparative analyses of genome architecture and show a considerable extent of variation in sequence composition during the recent evolution of *Zymoseptoria* lineages. We show that genome compartmentalization and accessory chromosomes represent shared ancestral traits among the pathogen species.

We identify extensive presence-absence variation of protein coding genes in genomes of five *Zymoseptoria* species consistent with the variable gene repertoire already reported for *Z. tritici* [28], Furthermore, the different species, share a particular genomic architecture that comprises specific accessory genome compartments. In spite of this variation, we observe an overall conserved synteny of the core chromosomes. In the *Zymoseptoria* genomes, we observe gene-dense, actively transcribed and H3K4me2-enriched compartments covering most of the core chromosomes. These compartments are clearly distinguishable from gene-sparse, non-transcribed and H3K4me2-deprived compartments. Based on previous analyses of accessory chromosomes in *Z. tritici*, we here consider this pattern as a specific hallmark of accessory genome compartments in the genus *Zymoseptoria* and not only in *Z. tritici* [6]. We hypothesize that these compartments likely represent accessory chromosomes in the different *Zymoseptoria* species.

We also identify accessory signatures in core chromosomes, including the previously described right arm of chromosome 7 [6]. Although this region has not been reported to share the same heavy pattern of presence-absence polymorphism as the accessory chromosomes, it is worth noting that a recent study identified a large deletion in chromosome 7 likely corresponding to this region in a single *Z. tritici* isolate originating from Yemen [37]. Here we show that this homologous region also exhibits accessory compartment hallmarks in the other *Zymoseptoria* species. A fusion between a core and an accessory chromosome occurred tens of thousands of years ago and prior to species divergence. The specific genomic and epigenetic features have remained stable through speciation and evolutionary time.

In this study, we confirm previously reported genome comparisons showing that gene content in *Z. tritici* is highly variable [28]. We further extended the identification of orthologs throughout the whole *Zymoseptoria* genus. Thereby, we show that more than 25% of the genes identified as *isolate-specific* in a comparison including only *Z. tritici* isolates are actually shared with its wild-grass infecting sister species, proving that a large proportion of the accessory genome of *Z. tritici* is not specific to this species. Instead, the accessory genome content is shared among *Zymoseptoria* species. The proportion of accessory *Z. tritici* genes shared with other *Zymoseptoria* species was found to be the highest in the Iranian isolate, which is the only isolate sympatric with the four sister-species. A likely explanation for this observation would be inter-specific gene flow which would allow the different wild species to exchange genes with sympatric *Z. tritici* isolates. This new finding is consistent with recent findings from population genomic data studies revealing extensive introgression between *Zymoseptoria* species [38, 39]. It also opens new perspectives for further analysis to understand how inter-specific gene flow has affected the evolution of the accessory genome of *Z. tritici*.

The genes with predicted functions and, in particular, functions related to pathogenicity are largely shared in *Zymoseptoria* genus. Although the lifestyles of the wild-grass infecting *Zymoseptoria* are poorly understood the species most likely share major features of their lifestyles. Thus, as expected, a similar CAZymes and BGC contents in all studied *Zymoseptoria* species was identified. Effector prediction is often used to better understand the adaptation of pathogens to their hosts [40]. However, in *Zymoseptoria*, the effectors are significantly enriched, not among the isolate- or even species-specific genes, but among the genes specific to the genus. If we exclude *Z. ardabiliae*, the repertoire of effectors shared at the genus level represents three-quarters of all predicted effectors. This ratio suggests either that only a small number of effectors confer host specificity among the different *Zymoseptoria* species or that host-specificity is conferred not by presence-absence of these effectors but by regulation of their expression [30]. In the *Botrytis* genus (Dothideomycetes), sister species infecting different hosts share effectors with confirmed functions [41]. We hypothesize that the different specificity levels reflect functional differences in the effector repertoire of *Zymoseptoria*. Effectors conserved across the *Zymoseptoria* genus are core effectors, potentially targeting key plant defense mechanisms common to all of their hosts [42]. In contrast, effectors which are *species-specific* or *isolate-specific* might reflect host specificity and target specific pathogen recognition pathways [42]. This assumption is supported by the fact that only a fraction of the *genus-specific* effectors is actually shared among all *Zymoseptoria* species while the majority shows presence-absence polymorphisms. *Z. ardabiliae* has been isolated from leaves of distantly related grass species in Iran, including *Lolium* spp., *Elymus repens,* and *Dactylis glomerata* and potentially resulting in a broader *species-specific* effector repertoire of *Z. ardabiliae* compared to the other species [21].

Consistent with a previous study [28], we found that *Z. tritici* core genes are more expressed compared to accessory genes *in planta*. Core genes are more likely to hold essential functions which could explain higher expression pattern during infection. Differentially expressed genes that are specifically induced during the course of the infection are very likely to have functions essential to pathogenicity of the fungus. Interestingly, here we show that within the differentially expressed genes between the biotrophic and the necrotrophic phases of infection, *isolate-* and *species-specific* genes have higher expression levels than core genes. These *isolate-* and *species-specific* genes could be functionally important and regulate functions linked to infection success in the biotrophic phase or to leaf colonization in the necrotrophic phase. Since these genes show presence-absence polymorphisms in the genus and in the *Z. tritici* species, they could represent a reservoir for possible adaptations to either host species, host cultivars or local environments.

## Conclusion

We investigated the genomic architecture in a genus of plant pathogens, including the economically relevant wheat pathogen *Z. tritici.* Comparing genome content and genome structure, we identified a large shared effector repertoire characterized by inter- and intraspecies presence-absence polymorphisms. Major features of genomic, transcriptomic and epigenetic compartmentalization, distinguishing accessory and core compartments, were shared among wheat and wild-grass infecting *Zymoseptoria* species. We conclude that compartmentalization of genomes is an ancestral trait in the *Zymoseptoria* genus.

## Methods and Materials

### Fungal material, DNA extraction and sequencing

Details regarding the individual *Zymoseptoria* isolates can be found in Table 1. For genomic data we used the three *Z. tritici* isolates IPO323 (reference), Zt05 and Zt10, one *Z. ardabiliae* isolate (Za17), one *Z. brevis* isolate (Zb87), one *Z. passerinii* isolate (Zpa63), and the *Z. pseudotritici* isolate (Zp13).For transcriptomic and epigenomic data we used *Z. tritici* Zt09 (IPO323 ΔChr18) a derivate of the reference isolate IPO323 deleted with the chromosome 18 [31].

Long read assemblies of the *Z. tritici* isolates Zt05 and Zt10 were described and published previously [30]. For DNA extraction and long read sequencing cultures of *Z. pseudotritici*, *Z. ardabiliae, Z. brevis* and *Z. passerinii* were maintained in liquid YMS medium (4 g/L yeast extract, 4 g/L malt extract, 4 g/L sucrose) at 200 rpm and 18°C. DNA extraction was conducted as previously described [26]. PacBio SMRTbell libraries were prepared using DNA extracted from single cells based on a CTAB extraction protocol [43]. The libraries were size selected with an 8-kb cutoff on a BluePippin system (Sage Science).

After selection, the average fragment length was 15 kb. Sequencing of the isolates Za17, Zb87, and Zp13 was run on a PacBio RS II instrument at the Functional Genomics Center, Zurich, Switzerland. Sequencing of the Zpa63 isolate was performed at the Max Planck-Genome-Centre, Cologne, Germany.

### Genome assembly, and repeat and gene predictions

For each isolate, we assembled the genome de novo using SMRT Analysis software v.5 (Pacific Bioscience) with two sets of parameters: default parameters and “fungal” parameters. We chose the best assemblies generated by comparison of all assembly statistics produced by the software Quast such as the number of finished contigs, the size of the assembly and the N50 [44]. Summary statistics for each assembly can be found in Table 1. In order to exclude poor quality contigs from the raw assemblies, we filtered out the contigs with less than 1.5X and more than 2X median read coverage as these might be unreliable from lack of data or because they contain only repeated DNA. This filter removed a high number of contigs, i.e., between 58% and 17% of contigs. Telomeric repeats (“CCCTAA”) were identified in the remaining contigs using bowtie2 to identify the number of fully assembled chromosomes or chromosomes arms based on the presence of more than six repeats at the contig extremities [45–47].

We next used the REPET package to annotate the repeat regions of *Z. ardabiliae* Za17, *Z. brevis* Zb87, *Z. pseudotritici* Zp13, *Z. passerinii* Zpa63, and the three *Z. tritici* isolates IPO323, Zt5 and Zt10 (https://urgi.versailles.inra.fr/Tools/REPET; [48, 49]. For each genome, we annotated the repetitive regions as follows: we first identified repetitive elements in each genome using TEdenovo following the developer’s recommendations and default parameters. The library of identified consensus repeats was then used to annotate the respective genomes using TEannot with default parameters.

We used several different approaches to predict gene coordinates and leveraged the information contained in previously published RNA sequencing data to increase the quality of the prediction [22, 30, 32]. We used GeneMark-ES for the first prediction *ab initio* using the option “--fungus” [50]. We furthermore used previously obtained RNAseq data in two different ways: first by mapping the reads to the respective genomes and second by assembling them *de novo* into longer transcripts. For this, we first trimmed the reads using Trimmomatic [51], and then mapped them on the newly assembled genomes using hisat2 [47]. Next we used the BRAKER1 pipeline to predict genes for each genome using the fungus flag and based on the previously mapped reads [52]. BRAKER applies GeneMark-ET and Augustus to create the first step of gene predictions based on spliced alignments and to produce a final gene prediction based on the best prediction of the first set [53, 54]. In addition to the *ab initio* gene predictions, this produced a second set of predicted genes. Furthermore, the RNAseq reads were separately assembled into gene transcripts using Trinity [55]. These were aligned using PASA and EVidence Modeler to produce consensus gene models from the two independent predictions and the *de novo* assembled transcripts [56]. Gene counts, length and other summary statistics presented in Table S1 were obtained using GenomeTools [57] and customs scripts.

The predicted gene sequences were the basis for an evaluation of the completeness of the assembly and gene prediction by the program BUSCO v.3 [58]. We used this method with the lineage dataset *Pezizomycotina.* The predicted genes were also used to create a phylogeny with the online implementation of CVtree3, using kmer sizes of 6 and 7 as recommend for fungi [59]. We generated a second tree with the whole assemblies, estimating a distance matrix using the andi software [60].

We predicted orthologs between the newly assembled genomes, the reference *Z. tritici* genome and the reference *Cercospora beticola* genome, a related Dothideomycete, which we used as outgroup to identify genes with orthologs restricted to the *Zymoseptoria* genus [21]. For this, we used the software PoFF [35, 61] which takes into account synteny information in the analyses of similarity inferred by the program Proteinortho [35].These orthogroups were used to visualize synteny between genomes using Circos [62].

The whole-genome assemblies were used to create a matrix distance with the software andi, from which we generated a tree [60]. A second tree was generated from the gene prediction with the online implementation of CVtree3 [59].

### Functional annotations

We used several tools to predict the putative functions for the gene models. First, we used the eggnog-mapper which provide COG, GO and KEGG annotations [63]. The online resource dbCAN2 was run to identify carbohydrate-active enzymes (CAZymes) [64]. Finally, for each genome, we used Antismash v3 (fungal version) to detect biosynthetic gene clusters (Figure S1; [65].

Additionally, we designed a pipeline to predict protein cell localization and to identify effector candidates. The pipeline for effector prediction is outlined in Figure S1 and includes the software DeepLoc [66], SignalP [67], TargetP [68], phobius [69] and TMHMM [70, 71], which predict the cellular location, the peptide signals and whether proteins are transmembrane. Effectors were identified with EffectorP v2 which uses both a new machine learning approach and more complete databases to improve effector prediction compared to the previous version [19]. The pipeline also includes software which are specifically targeted to annotate plant pathogenic functions, namely the program ApoplastP [72] and LOCALIZER [73]. We wrote wrappers scripts, which run the software and create consensus between the different prediction tools providing one command line from the user. These scripts are available at https://gitlab.gwdg.de/alice.feurtey/genome_architecture_zymoseptoria. Briefly, we gathered outputs of several software to predict the cellular location, transmembrane domain and secretion and created a consensus based on the different output to prevent the pitfalls of any one of these methods. From this consensus, we extracted the gene products predicted to be secreted and without a transmembrane domain. The comparisons of genes functions repartition were done by combining predictions of COG categories, secondary metabolite genes with pathogenicity-related gene functional categories such as CAZymes and effector predictions.

### Gene expression analyses

To update expression profiles on the new genome assemblies and new gene predictions of the three *Z. tritici* isolates, we used previously generated RNA-seq data from *in planta* and *in vitro* growth [30, 32]. The *in planta* RNAseq data was obtained from infected leaves at four different stages corresponding to early and late biotrophic (stage A and stage B) and necrotrophic (stage C and stage D) stages of the three *Z. tritici* isolates [30]. Strand-specific RNA-libraries were sequenced using Illumina HISeq2500, with 100pb single-end reads for a total read number ranging from 89.5 to 147.5 million reads per sample. This data was previously analyzed [30], using gene predictions generated from an Illumina-based assembly [22]. The reads were here mapped on the new assemblies of Zt5 and Zt10 and the reference genome of IPO323 after trimming. We used the DESeq2 R package to determine differential gene expression during *in planta* infection, considering only two infection stages; biotrophic and necrotrophic [74]. Gene expression was assessed as Transcript per Million (TPM). Briefly, TPM is calculated by normalizing read counts with coding region length resulting in the number of reads per kilobase (RPK). RPK total counts per sample are then divided by 1 million to generate a “per million” scaling factor. We calculated the coding region length of each gene with GenomicFeatures R package using the function called “exonsBy” [75]. For gene expression analyses, we further filtered our gene predictions to remove any predicted transposases and other TE-related annotations based on the Eggnog mapper annotations.

### ChIP-sequencing and data analysis

*Z. ardabiliae* (Za17) and *Z. pseudotritici* (Zp13) cells were grown in liquid YMS medium for two days at 18°C until an OD_600_ of ~ 1 was reached. Chromatin immunoprecipitation and library preparation were performed as previously described [76]. We sequenced two biological and two technical replicates per isolate and used antibodies against the euchromatin histone mark H3K4me2 (#07–030, Merck Millipore). Sequencing was performed at the OSU Center for Genome Research and Biocomputing (Oregon State University, Corvallis, USA) on an Illumina HiSeq2000 to obtain 50-nt reads. The data was quality-filtered using the FastX toolkit (http://hannonlab.cshl.edu/fastx_toolkit/), mapping was performed using bowtie2 [77] and peaks were called using HOMER [78]. Peaks were called individually for each replicate, but only peaks that were detected in all replicates were considered and merged for further analysis. Merging of peaks and genome wide sequence coverage with enriched regions was assessed using bedtools [46].

## Data availability

The assembled genomes can be found at 10.5281/zenodo.3568213. The gene annotations are deposited at 10.5281/zenodo.3568213. The functional annotation pipeline, additional scripts and command lines used to create the results presented in this manuscript can be found at https://gitlab.gwdg.de/alice.feurtey/genome_architecture_zymoseptoria.

## Ethics approval and consent to participate

Not applicable.

## Consent for publication

Not applicable.

## Availability of data and materials

All the data supporting the findings of this study are openly available at 10.5281/zenodo.3568213.

## Competing interests

The authors declare that they have no competing interests.

## Funding and Acknowledgements

CL is funded by the Institut national de la recherche agronomique (INRA) in the framework of a “Contrat Jeune Scientifique” and by the Labex ARBRE (Lab of Excellence ARBRE). Genome research in the group of EHS is funded by the DFG Priority Program SPP1819 and the Canadian Institute for Advanced Research (CIFAR). AF is funded by by the DFG Priority Program SPP1819.

## Authors’ contributions

AF, CL and MM performed data and results analyses. EHS, AF and CL contributed to the design and implementation of the research. AF, CL, CE and DC performed genome assemblies. MH, JH, MM, MF and KS performed the experimental procedures. All authors contributed to the writing of the manuscript.

## Supplementary Tables

Table S1: Summary statistics from gene predictions generated in this study.

Table S2: Summary of gene expression in *Z. tritici* isolates during infection stages

Table S3: Enrichment of species-specific and isolate-specific genes per chromosome/contig

## Supplementary Figure Legends

**Figure S1:** Plant-associated genes compartmentalization along the chromosomes. The first track represents core (dark grey) and accessory (light grey) chromosomes/contigs. Fully assembled contigs are marked by the presence of telomeric repeats (orange). Circles from outside to inside represent the position of: effector genes (blue), biosynthetic gene clusters (BGC, green) and CAZymes (yellow), respectively.

**Figure S2:** A) Intra-species synteny between the reference genome of *Z. tritici* and the genome of the *Z. tritici* isolate Zt05. Each color represents a different chromosome as based on the reference *Z. tritici* genome, except for accessory chromosomes, which are in grey. The connecting lines represent orthologs between each genome. The track between the chromosomes and connecting lines are predicted effectors. B) Inter-species synteny between reference genome of *Z. tritici* and *Z. pseudotritici*. The arrows represent the large-scale inversions identified between the genomes of these two species.

**Figure S3:** Inter-species synteny between genomes of *Z. pseudotritici* (dark grey) and *Z. brevis* (blue). The arrows indicate the contigs affected by the large-scale inversions identified between *Z. tritici* and both *Z.pseudotritici* and *Z. brevis*.

**Figure S4:** Genome architecture in A) *Z. tritici* Zt05, B) *Z. tritici* Zt10. C) *Z. brevis* Zb87 and D) *Z. ardabiliae* Za17. Circles from the outside to the inside represent respectively: gene density along chromosomes/contigs; gene expression in vitro (TPM); H3K4me2 distribution in vitro (only for Za17) and species-specific gene density

**Figure S5:** Simplified diagram of the pipeline used to predict the functions and subcellular localization of gene model products.

**Figure S6:** Functional gene categories in *Zymoseptoria spp*. genomes. A) The number of genes in Eggmapper COG categories in addition to Effectors and CAZymes. B) Pathogenicity-related genes of interest: secreted proteins, effectors, secreted CAZymes and non-secreted CAZymes. C) Subcellular localization of predicted gene products.

**Figure S7:** Principal component analysis of DESeq2 rlog transformed expression data.

